# Validation and characterization of a DNA methylation alcohol biomarker across the life course

**DOI:** 10.1101/591404

**Authors:** Paul Darius Yousefi, Rebecca Richmond, Ryan Langdon, Andrew Ness, Chunyu Liu, Daniel Levy, Caroline Relton, Matthew Suderman, Luisa Zuccolo

**Author notes:** Corresponding author: Luisa Zuccolo, Oakfield House, Oakfield Grove, Bristol, BS82BN UK.

## Abstract

Recently, an alcohol predictor was developed using DNA methylation at 144 CpG sites (DNAm-Alc) as a biomarker for improved clinical or epidemiologic assessment of alcohol-related ill health. We validate the performance and characterize the drivers of this DNAm-Alc for the first time in independent populations. In N=1,049 parents from the Avon Longitudinal Study of Parents and Children (ALSPAC) Accessible Resource for Integrated Epigenomic Studies (ARIES) at midlife, we found DNAm-Alc explained 7.6% of the variation in alcohol intake, roughly half of what had been reported previously, and interestingly explained a larger 9.8% of AUDIT score, a scale of alcohol use disorder. Explanatory capacity in participants from the offspring generation of ARIES measured during adolescence was much lower. However, DNAm-Alc explained 14.3% of the variation in replication using the Head and Neck 5000 (HN5000) clinical cohort that had higher average alcohol consumption. To investigate whether this relationship was being driven by genetic and/or earlier environment confounding we examined how earlier vs. concurrent DNAm-Alc measures predicted AUDIT scores. In both ARIES parental and offspring generations, we observed associations between AUDIT and concurrent, but not earlier DNAm-Alc, suggesting independence from genetic and stable environmental contributions. The stronger relationship between DNAm-Alcs and AUDIT in parents at midlife compared to adolescents despite similar levels of consumption suggests that DNAm-Alc likely reflects long-term patterns of alcohol abuse. Such biomarkers may have potential applications for biomonitoring and risk prediction, especially in cases where reporting bias is a concern.

## Introduction

Alcohol use and misuse are responsible for a large proportion of the global burden of disease,^1^ however measuring alcohol exposure in epidemiological studies presents a number of challenges. Self-reported alcohol intake is the most commonly used source of information, but such measurements are fraught with both error and bias, and generally underestimate exposure.^2,3^ The latter is often more pronounced at higher levels of exposure as heavy drinkers can lose track of their intake or differentially under-report their consumption compared to light drinkers, which may result in an overestimation of the effects of alcohol and an underestimation of the burden of disease. Similar bias can occur in retrospective studies where the quality of self-reported alcohol use can vary between cases and controls.

Objective biomarkers of alcohol intake are useful tools for epidemiology and public health because they have the potential to improve exposure assessment, especially when reporting bias may be a major issue, and inform screening and risk prediction strategies. Currently, alcohol biomarkers exist for exposure assessment over short-term windows (e.g. breathalysers in toxicology) or at high doses (e.g. liver enzymes and other blood-based laboratory markers for excessive alcohol use).^4^ What is lacking is a reliable biomarker reflecting the distribution of alcohol intake across the range observed in the general population and that is capable of capturing exposure over months and years.

DNA methylation, currently the best characterized epigenetic modification, which is involved in regulating gene expression, has been found to be altered in relation to several environmental and biological factors.^5–9^ Patterns of DNA methylation represent attractive targets for developing novel biomarkers of exposure as studies have shown that DNA methylation levels at small numbers of or even individual CpG sites in non-invasive peripheral tissues can serve as sensitive and specific indicators of exposure.^10,11^ Further, DNA methylation signals are stable over highly variable timescales, with some having been observed to persist well beyond the half-lives of existing targets for alcohol biomarkers.^12^ For example, associations observed in relation to prenatal smoking exposure have been seen to endure into childhood^13^ and some effect of the exposure appears detectable into adulthood.^14^

Liu et al. developed four new blood DNA methylation-based alcohol models (DNAm-Alcs) as biomarkers of alcohol intake, using a large population sample (the CHARGE consortium^15^). Of these four models, the one comprising the highest number of CpG sites (144) consistently showed the best performance.^6^ The well powered analyses, good range of alcohol intake and promising proportion of the variance explained make this an interesting candidate biomarker. However, the reported performance of this model is likely inflated due to over-fitting.^16^ Another limitation is the use of cross-sectional data makes it difficult to determine the direction of the association or to rule out genetic confounding, which could be an issue since many CpGs having been shown to be under genetic control^17^ and alcohol use disorders are heritable.^18^

Here and for the first time, we aim to assess the out-of-sample performance of the DNAm-Alcs by using data at multiple time points from an independent population, the subset of participants from the Avon Longitudinal Study of Parents and Children (ALSPAC) included in the Accessible Resource for Integrated Epigenomic Studies (ARIES), and further replication using the Head and Neck 5000 (HN5000) clinical cohort. Repeated measures of alcohol use and blood DNA methylation in two generations of the ARIES study and at different points in the life course, including DNA methylation profiles collected prior to own alcohol exposure, were used to address our aims. Our first aim was to test the performance of the DNAm-Alcs in ARIES and HN5000. Second, we aimed to further investigate the performance of DNAm-Alcs in ARIES at predicting measures of alcohol use disorders (AUDIT scores) in different groups of individuals (males and females at midlife and adolescence), which is designed to estimate the extent to which an individual engages in harmful patterns of alcohol consumption.^19^ Lastly, we sought to address additional details regarding the nature of the association between the DNAm-Alcs and alcohol use, such as duration of alcohol use and the potential for genetic confounding, by using measurements of alcohol and DNA methylation at different times in the life course.

## Materials and methods

### ARIES study

The Avon Longitudinal Study of Parents and Children (ALSPAC) is a population-based birth cohort drawn from the South West of England. Pregnant women resident in Avon, UK with expected dates of delivery 1st April 1991 to 31st December 1992 were invited to take part in the study. The initial number of pregnancies enrolled is 14,541 (for these at least one questionnaire has been returned or a “Children in Focus” clinic had been attended by 19/07/99). Of these initial pregnancies, there was a total of 14,676 foetuses, resulting in 14,062 live births and 13,988 children who were alive at 1 year of age. Detailed information on the mothers and their offspring has since been regularly collected.^20,21^ The study website contains details of all the data that is available through a fully searchable data dictionary and variable search tool (http://www.bristol.ac.uk/alspac/researchers/our-data/). Ethical approval for the study was obtained from the ALSPAC Ethics and Law Committee and the Local Research Ethics Committees (http://www.bristol.ac.uk/alspac/researchers/research-ethics/). Informed consent for the use of data collected via questionnaires and clinics was obtained from participants following the recommendations of the ALSPAC Ethics and Law Committee at the time. Consent for biological samples has been collected in accordance with the Human Tissue Act (2004).

Blood from 1 018 mother–child pairs (children at three time points and their mothers at two time points) were selected for analysis as part of the Accessible Resource for Integrative Epigenomic Studies (ARIES, http://www.ariesepigenomics.org.uk/).^22^ ARIES consists of DNA methylation profiles obtained from 1 018 mother-child pairs measured at five time points (three time points for children: birth, childhood and adolescence; and two for mothers: during pregnancy and at midlife), and on approximately 600 fathers at midlife. In this analysis, we included participants from the ARIES parental generation at midlife (N = 1 049, mean age = 50.2 ± 5.4 SD) and the offspring generation at adolescence (N = 626, mean age = 17.4 ± 0.9 SD) with DNA methylation, alcohol, and covariate information available (Figure 1). The subset of these participants with DNA methylation available at an earlier time point, during pregnancy in the parents (N = 518, mean age = 29.1 ± 4.2 SD) and in cord blood at birth (N = 438) and childhood (N = 601, mean age = 7.4 ± 0.1 SD) in the offspring, were included in some analyses.

**Figure 1.**
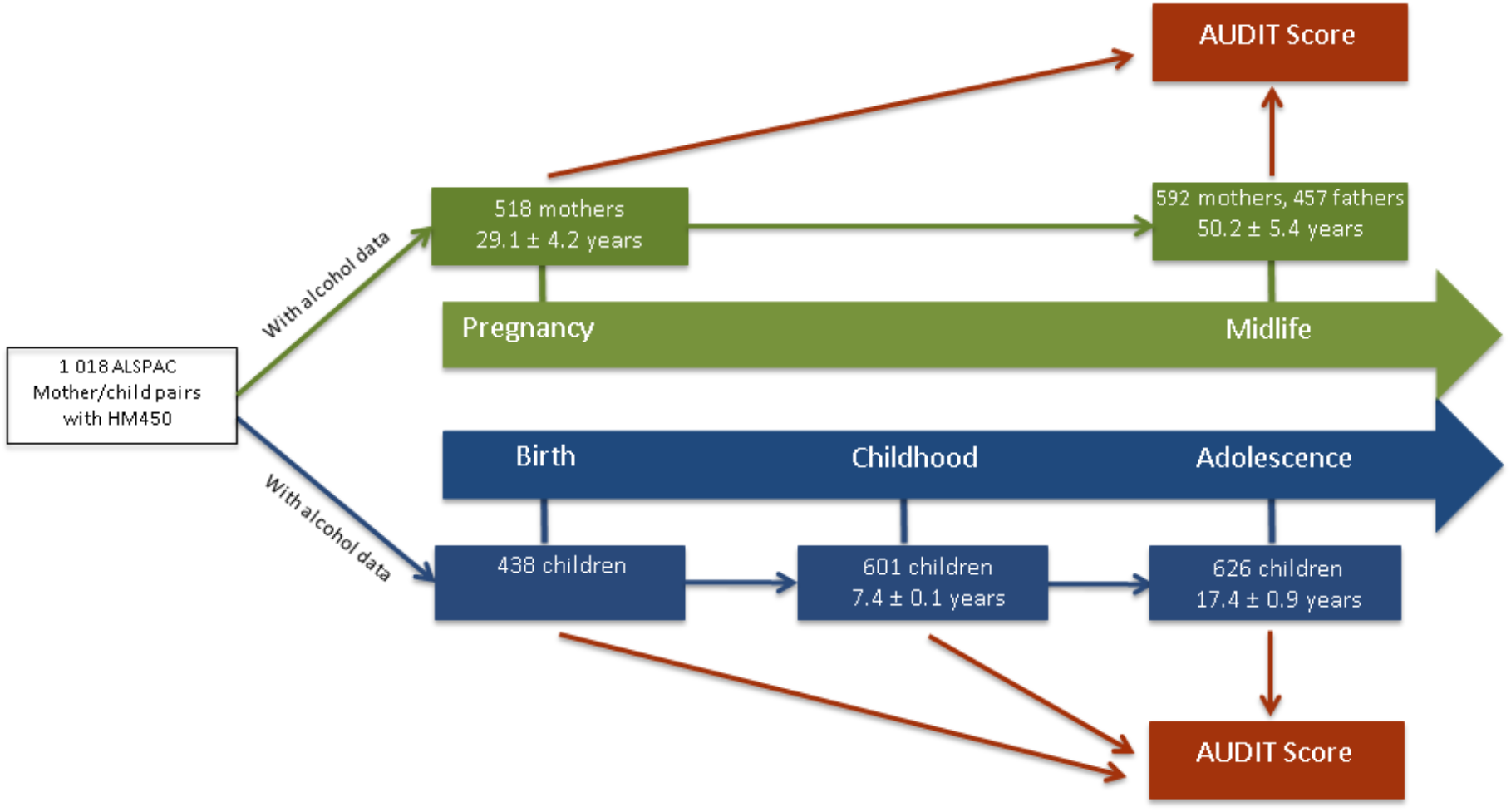
Flow chart of participants within ARIES with HM450 and alcohol questionnaire data available by time-point of collection and schematic of early vs. cross-sectional analysis performed. Abbreviations: ARIES, Accessible Resource for Integrated Epigenomic Studies

### HN5000 replication study

The Head and Neck 5000 (HN5000) clinical cohort study, a clinical cohort study of individuals with head and neck cancer, was used as an additional independent replication population. Full details of the study methods and overall population are described in detail elsewhere.^23,24^ Briefly, between April 2011 and December 2014, 5511 individuals with HNC were recruited from 76 centres across the UK. Individuals were recruited before they started treatment unless their treatment was their diagnostic procedure. Full ethical approval was granted by the South West Frenchay Regional Ethics Committee (ref: 10/H0107/57).

At baseline, participants were asked to complete a series of questionnaires and information on diagnosis, treatment and co-morbidity was recorded on a short data capture form. Diagnoses were coded using the International Classification of Diseases (ICD) version 10^25^ and clinical staging of the tumour was derived based on the American Head and Neck Society TNM staging.^26^ Biological samples (blood, saliva and tissue) were collected from all consenting participants, and DNA methylation was measured for 448 individuals with an ICD-10 coding of oropharynx cancer and complete baseline questionnaire and data capture information.^23^ Of those, participants of European ancestry with DNA methylation, alcohol, and all covariate information available were included in this analysis (N=281, mean age = 58.4 ± 9.9 SD)

### Alcohol intake and disorder

Alcohol intake and AUDIT score^27^ were available at multiple time points for the ALSPAC participants. To validate the performance of the DNAm-Alcs, we utilised measures of alcohol intake at midlife and adolescence taken as close as possible to the DNAm measurement, to replicate the cross-sectional design used in the derivation and original evaluation of DNAm-Alcs. For ARIES parents at midlife and the offspring generation at adolescence, alcohol drinking frequency was assessed by response to the question, “How often do you have a drink containing alcohol”, with possible responses including “Never”; “Monthly or less”; “2 to 4 times a month”; “2 to 4 times a week”; “4 or more times a week”. “Never” respondents were considered non-drinkers and were excluded from all analyses apart from examination of their influence in sensitivity analysis.

Participants were also asked to report the quantity of their typical alcohol consumption where “one drink referred to ½pint of beer/cider, a small (125ml) glass of wine or a single (25ml) measure of spirit”, each of which is roughly equivalent to one UK alcohol unit (8 grams of ethanol). Alcohol intake was then calculated by multiplying participant drinking frequency by the typical quantity consumed and converting from UK units/week to g/day. In all statistical models, alcohol intake was log-transformed (log (g/day + 1)) to coerce its distribution to approximate normality. For categorical analyses, drinkers were grouped according to the following thresholds of alcohol intake: ‘non-drinkers’ reported no alcohol consumption (intake g/d = 0), ‘light drinkers’ consumed 0 < g/day ⩽14 for women and 0 < g/day ⩽28 in men, ‘heavy drinkers’ ⩾28 g/day in men and ⩾42 g/day in men. These categorical thresholds, which are the same as those used by Liu et al., were selected in order to maximize comparability between our results.

Alcohol use disorder identification test (AUDIT) scores were taken from the same time points as the measures of alcohol intake, cross-sectionally to DNAm at midlife in the ARIES parental generation and in adolescence in the offspring. AUDIT includes responses to 10 questions that evaluate participant alcohol misuse in the past year, producing scores that range from 0 to 40. Self-reported non-drinkers are included in this scale, receiving a score of 0.

In HN5000, detailed information on alcohol history was obtained at baseline via the self-reported questionnaire. Participants were asked about their current drinking status and their use of alcohol prior to receiving their HNC diagnosis. Respondents were asked to report their average weekly alcohol consumption of a range of beverage types (wine, spirits, and beer/larger/cider) before they were diagnosed with cancer. From these measures, average intake of alcohol consumption in units per week was derived, which we subsequently converted to log (g/day + 1).

### DNA methylation measurements

In ARIES, cord blood and peripheral blood samples (whole blood, buffy coats or blood spots) were collected according to standard procedures. DNA extraction, wet laboratory preparation and DNA methylation measurement by Illumina Infinium HumanMethylation450K (HM450) BeadChip was performed as part of the ARIES project, as previously described.^22,28^ Briefly, samples from all ARIES time-points were distributed semirandomly across HM450 slides to minimize the potential for confounding by technical batch. Data pre-processing was performed using the *meffil* package.^29^ Samples failing quality control (average probe detection P value ≥0.01, those with sex or genotype mismatches) were excluded from further analysis, and probes containing <95% of signals detectable above background signal (detection P value <0.01) were also removed. Functional normalization^30^ was performed to minimize non-biological variation in probes.

In HN5000, blood samples were processed and frozen upon receipt and stored at - 80°C. Genome-wide DNA methylation data was generated in participants using Infinium MethylationEPIC BeadChip (EPIC array) (Illumina, USA) at the Bristol Bioresource Laboratories. Following extraction, DNA was bisulphite-converted using the Zymo EZ DNA MethylationTM kit (Zymo, Irvine, CA, USA) ang genome-wide methylation status of over 850,000 CpG sites was measured using the EPIC array according to the manufacturer protocol using an Illumina iScan. All data pre-processing and normalization was again performed using the *meffil* package,^29^ which excluded 5 of the total samples for not meeting the QC criteria.

### Covariates

For the ARIES parental generation at midlife, offspring generation in both childhood and adolescence, and HN5000 participants, DNA methylation levels were adjusted for age, sex, body mass index (BMI) and Houseman estimate white blood cell (WBC) counts by residualizing on these factors by ordinary least squares (OLS) regression prior to calculation of DNAm-Alcs.^31,32^ Levels from pregnancy were not adjusted for sex, as only female participants had DNA methylation available, but were adjusted for age, WBC counts and prepregnancy BMI. The adjustment set for DNA methylation at birth included gestational age (weeks), child sex, birthweight (g), and WBC counts calculated using a cord blood reference set.^33^ Indicator variables for self-reported current smoking at midlife for the ARIES parents and for having ever/never smoked a cigarette in adolescence for the offspring were utilised in sensitivity analyses.

### DNA methylation alcohol score

We computed DNA methylation-derived alcohol scores (DNAm-Alcs) using coefficients made available from the lasso models estimated by Liu and colleagues,^6^ predicting the relationship between DNA methylation and alcohol intake. Four DNAm-Alcs were examined using different sets of coefficients that varied according to the lasso tuning parameters considered by the original authors and the number of CpG sites they utilised as inputs (5, 23, 78, and 144 CpGs). All CpG sites with coefficients used in the smaller DNAm-Alcs were subsets of the CpGs used in the DNAm-Alcs with larger sets of coefficients. DNAm-Alc values were computed as weighted scores by multiplying each set of Liu et al. coefficients against the DNA methylation levels of ARIES and HN5000 participants at the corresponding CpG sites, and summing these values:

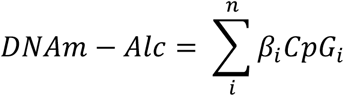

DNA methylation levels were residualized from the linear effects of the covariates described above prior to computing DNAm-Alc. Software for producing all four DNAm-Alc estimates is freely available using the *dnamalci* R package (https://github.com/yousefi138/dnamalci).

### Statistical analysis

Overall performance of DNAm-Alc scores in predicting continuous measures, including alcohol intake and AUDIT score, was assessed by R^2^ or the percent of the variance in the outcome variable explained. For binary outcome measures, receiver operating characteristic (ROC) curves were generated and area under the curve (AUC) was calculated using the pROC package^34^ to evaluate predictive performance. Linear relationships between DNAm-Alcs and AUDIT scores from different time points were evaluated using OLS regression. Models involving differing numbers of explanatory parameters were compared using the OLS-derived adjusted R^2^ in order to account for the discrepancy in model degrees of freedom.

We further performed sensitivity analyses to check the robustness of our results to the impact of smoking and the inclusion of non-drinkers. To this end, we ran cross-sectional models using DNAm values that had been residualised using concurrent smoking as an additional covariate, and excluding participants identified as self-reported non-drinkers.

As an additional comparison, we re-fit linear models with the same sets of CpGs used in each DNAm-Alc, as well as age, sex, and BMI as explanatory variables. This provided an inflated assessment of prediction due to over-fitting, as has been described in detail by Hattab, Clark, and van den Oord,^16^ but allowed direct comparison to the performance of the models originally implemented by Liu et al. In these re-estimated models, adjusted R^2^ from the OLS model was again used to compare models with different numbers of covariates.

All statistical analysis was performed in R version 3.3.1.^35^

## Results

### Validation of DNAm-Alcs

The four DNAm-Alcs evaluated, using 5-144 CpGs, explained between 3.9 and 7.6% of the variation in alcohol intake in our sample of N=1 049 ARIES participants at midlife, using DNA methylation levels independent of age, sex, and BMI (Table 1; Figure 1). Results were identical when adjusting for concurrent smoking (3.8 - 7.6%; Supplementary Table 1) or excluding participants who had reported being non-drinkers (3.8 – 7.6%; Supplementary Table 2). Although mean self-reported alcohol intake was similar between ARIES adolescents and their parents at midlife (mean intake = 8.1 ± 10.2 and 8.2 ± 8.8 g/d respectively, Figure 2), DNAm-Alcs explained a much lower percentage of the variation in adolescent alcohol intake, with a maximum of 0.8% for the 144 CpG DNAm-Alc. Again, this result did not change substantially when accounting for smoking or when excluding non-drinkers.

**Figure 2.**
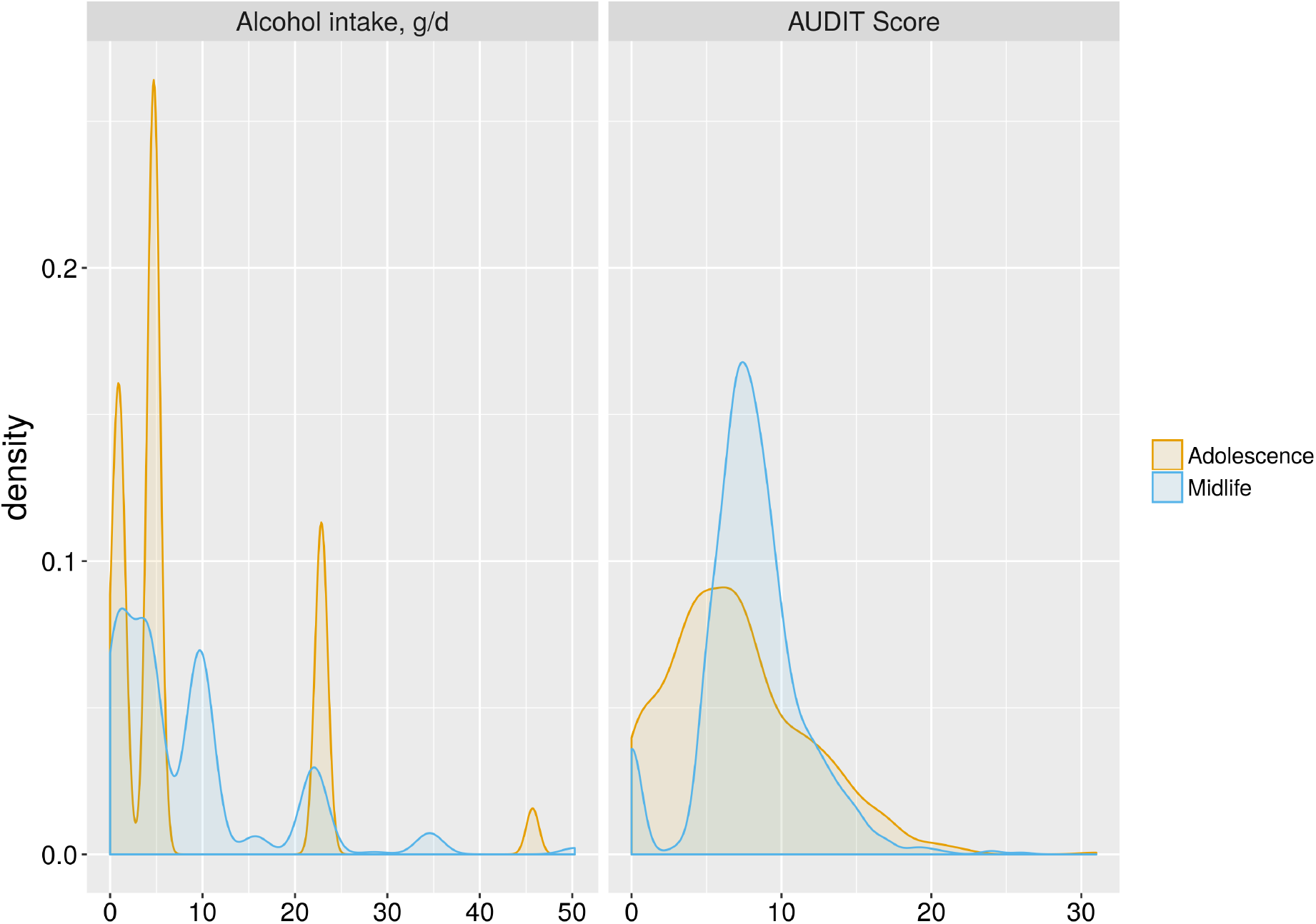
Distributions of alcohol use measurements in ARIES participants. Abbreviations: ARIES, Accessible Resource for Integrated Epigenomic Studies

**Table 1.**
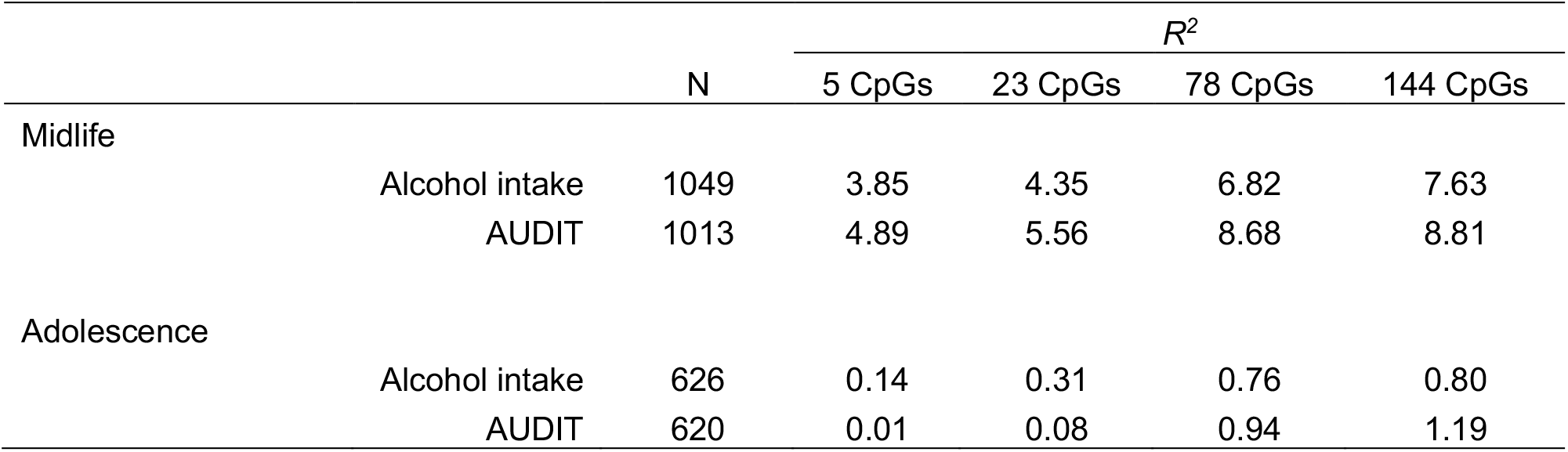
R^2^ between DNAm-Alcs and measures of alcohol use, alcohol intake (log(g/day + 1)) and AUDIT score, in ARIES parents at midlife and offspring at adolescence. Abbreviations: DNAm-Alcs, DNA methylation alcohol biomarkers; AUDIT, alcohol use disorders identification test; ARIES, Accessible Resource for Integrated Epigenomics Studies.

In each of these analyses, prediction performance was positively correlated with the number of CpG sites included in the model. That is, the 5 CpG DNAm-Alc consistently explained the least variation, the 144 CpG explained the most, with the 23 CpG and 78 CpG models ordered in between.

Replication in an independent UK-based population of adults with oropharyngeal cancer, N= 281 participants from HN5000, found each of the four DNAm-Alcs explained higher levels of variation in alcohol intake than observed in ARIES, with adjusted R^2^ ranging from 8.8% to 14.3% (Supplementary Table 3). While this population was similar in age to the ARIES participants at midlife (mean age = 58.4 ± 9.9), their levels of alcohol consumption were considerably higher (mean intake = 28.0 ± 31.3g/d).

Similar trends but with consistently stronger correlations were observed in ARIES parents at midlife when the coefficients of the CpG sites included in the DNAm-Alcs were re-estimated in ARIES, to allow direct comparison with performances based on re-estimated models reported by Liu et al. (Supplementary Table 4). By this approach, adjusted R^2^ ranged from 8.8 to 17.3% for each of four CpG covariate sets, including the independent explanatory effects of age, sex, and BMI that jointly contributed an adjusted R^2^ of 4.8%.

Assessment of the ability of DNAm-Alcs to distinguish heavy drinkers from nondrinkers and heavy drinkers from light drinkers by ROC analysis produced AUC values ranging from 0.48 to 0.57 and 0.55 to 0.57 respectively in ARIES adults at midlife (Figure 3). Prediction performance did not differ by the number of CpG sites included in DNAm-Alc: the 23 CpG DNAm-Alc produced the largest AUC, 0.57, in resolving heavy versus non-drinkers, and both the 78 and 144 CpG DNAm-Alcs had identical AUCs at predicting heavy versus light drinkers. In ARIES adolescents, AUCs ranged from 0.53 to 0.60 and 0.47 to 0.55 for heavy versus nondrinkers and light drinkers respectively, meaning some performed no better than chance alone (AUC = 0.50). While the score with the best predictive performance was the 144 CpG DNAm-Alc, with an AUC of 0.60 for heavy versus non-drinkers, there was again no trend observed in the predictive performance as CpG coefficients were added to the DNAm-Alcs, the same 144 CpG DNAm-Alc in fact performing the poorest in comparing heavy versus light drinkers (AUC < 0.50),.

**Figure 3.**
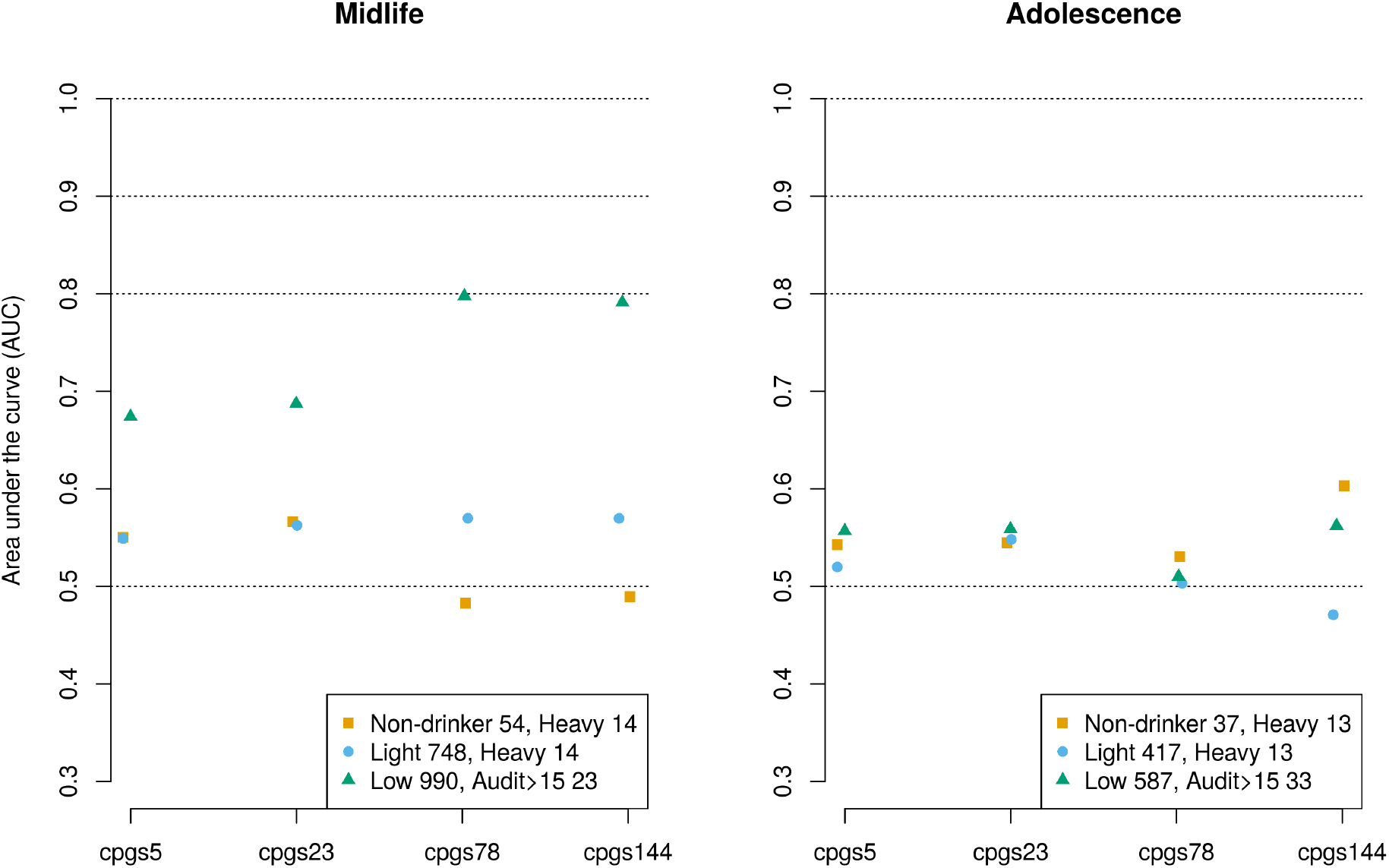
Area under the curve for prediction of binary alcohol categories by DNAm-Alcs. ROC analysis was performed to discriminate categories of alcohol intake (heavy versus nondrinkers and heavy versus light drinkers) and AUDIT score in ARIES parents at midlife (left figure) and children at 17 years of age (right figure). ‘Heavy drinkers’ were participants who consumed ≥ 42 g per day in men and ≥ 28 g per day in women; ‘non-drinkers’ consumed 0 g per day; ‘light drinkers’ consume 0 < g per day ≤ 28 in men and 0< g per day ≤ 14 in women; AUDIT was dichotomized into participants with a score >15 and those with a score ≤ 15. Abbreviations: DNAm-Alcs, DNA methylation alcohol biomarkers; AUDIT, alcohol use disorders identification test; ARIES, Accessible Resource for Integrated Epigenomics Studies.

### DNAm-Alc and AUDIT score

At midlife, DNAm-Alcs explained from 4.9 to 8.8% of the variation in participant AUDIT scores, a measure of alcohol use disorder (Table 1). In fact, every DNAm-Alc with a particular number of CpGs explained more of the variation in AUDIT than for alcohol intake. On average, DNAm-Alcs explained 1.3% more variation of AUDIT than alcohol intake in midlife. These DNAm-Alcs explained a comparable but slightly greater amount of variation in AUDIT when performing additional adjustment for smoking or following the exclusion of non-drinkers, from 5.0 to 9.0% and 5.5 to 9.8% respectively (Supplementary Tables 1 and 2).

In adolescents, DNAm-Alcs explained substantially less variation in AUDIT than at midlife, reaching only a maximum of 1.2% for the 144 CpG biomarker. Despite this decrease in total variance explained among adolescents, the DNAm-Alcs continued to correlate slightly better with AUDIT than with alcohol intake, explaining 0.05% more on average. Similar relationships with AUDIT were once again observed when adjusting for smoking and excluding non-drinkers.

As with alcohol intake, we observed a trend of consistent improvement in prediction performance for AUDIT with additional CpG sites: the 144 CpG DNAm-Alc score always performing the best and the 5 CpG DNAm-Alc performing worst.

The ability of the DNAm-Alcs to distinguish high (>15) versus low AUDIT score categories in ROC analysis produced AUCs that ranged from 0.67 with the 5 CpG DNAm-Alc to 0.80 for the 78 CpG score in ARIES participants at midlife (Figure 3). These AUC values were uniformly higher than those observed for all DNAm-Alcs at predicting heavy drinkers from light or non-drinkers using alcohol intake derived cut-offs. In ARIES adolescents, DNAm-Alc prediction of high versus low AUDIT individuals was much more comparable with performance observed using intake-based categories, with AUCs ranging from 0.51 to 0.56.

### Earlier vs. cross-sectional DNAm-Alc and AUDIT

To determine whether the relationship observed between DNAm-Alc and AUDIT using cross-sectional data could be explained by genetic confounding, we compared the predictive performance of DNAm-Alc based on methylation of blood drawn earlier in life, at a period of expected low or no alcohol use, to the predictive performance of DNAm-Alc based on methylation of blood drawn later in the same participants, at the same time as completion of the AUDIT questionnaire. If we show that DNAm-Alc from blood drawn earlier in life is not associated with later AUDIT, then we’ve ruled out both genetic and earlier environment confounding of the cross-sectional DNAm-Alc – AUDIT relationship. A schematic flow-chart describing this analysis is provided in Figure 1.

In the ARIES mothers, we compared the effects of the 144 CpG DNAm-Alc measured during pregnancy to those of DNAm-Alc measured cross-sectionally to AUDIT, 21.1 years later on average during midlife (Table 2). For the N=518 participants with DNAm-Alc at both time-points and AUDIT available, we found no evidence for a relationship between DNAm-Alc during pregnancy and AUDIT at midlife, both when considered as a single independent variable (β = 0.15, p = 0.56) and when adjusting for DNAm-Alc at midlife (β = 0.14, p = 0.57). Conversely, we observed a strong relationship between DNAm-Alc at midlife and AUDIT (β = 1.63, p < 0.001) even when accounting for pregnancy DNAm-Alc (β = 1.66, p < 0.001).

**Table 2.**
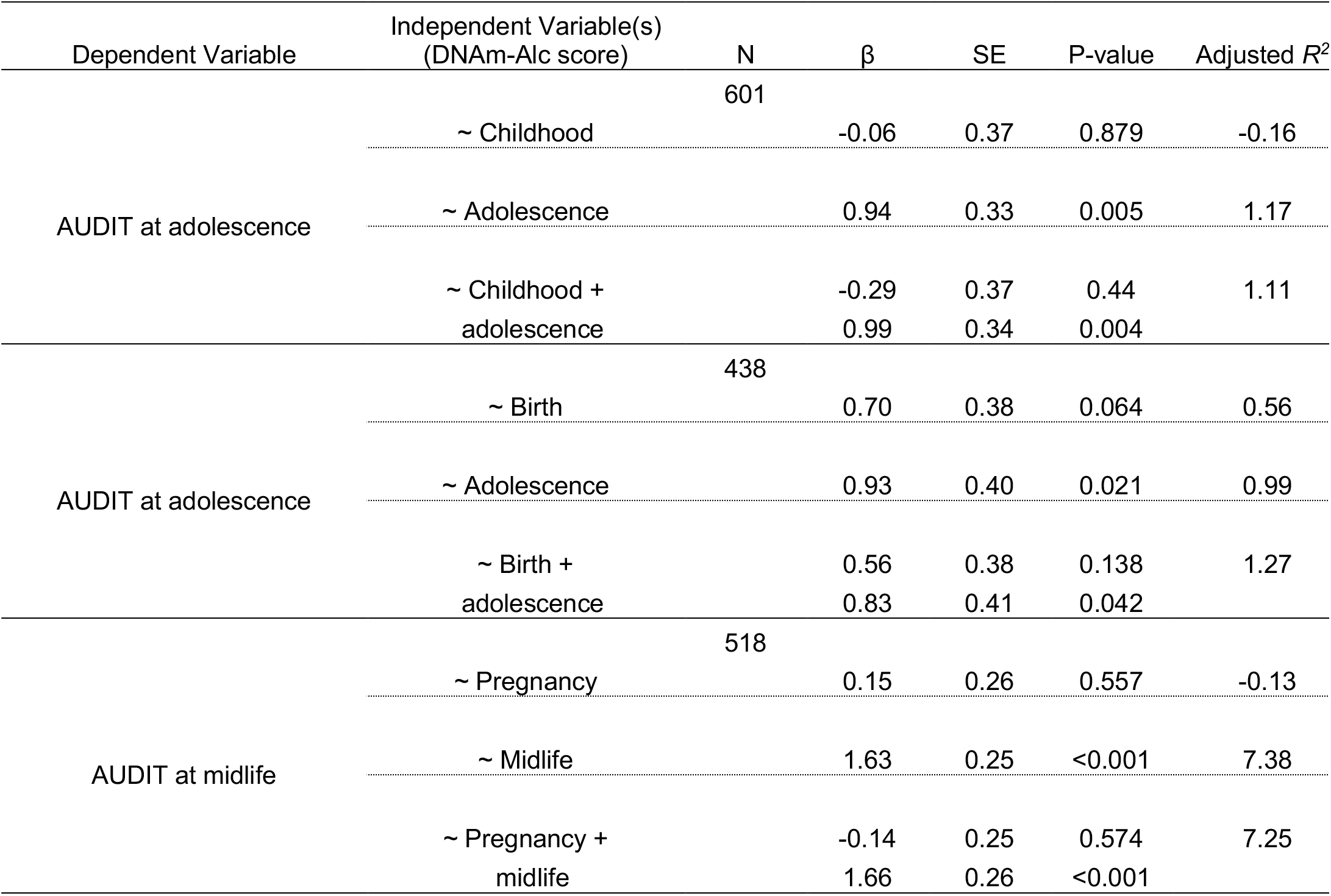
DNAm-Alc and AUDIT score in the sub-sample with repeated DNAm-Alc measurements. Estimates of the effect of DNAm-Alc on AUDIT at a time point prior to AUDIT measurement (childhood, birth, and pregnancy), at a time point cross-sectional to AUDIT (adolescence and midlife), and with the effect of prior and cross-sectional DNAm-Alc modelled simultaneously. *See Methods for details. Abbreviations: AUDIT, alcohol use disorders identification test; DNAm-Alcs, DNA methylation alcohol biomarkers.

| Dependent Variable | Independent Variable(s)<br>(DNAm-Alc score) | N | $\beta$ | SE | P-value | Adjusted $R^2$ |
| --- | --- | --- | --- | --- | --- | --- |
| AUDIT at adolescence |  | 601 |  |  |  |  |
|  | ~ Childhood |  | -0.06 | 0.37 | 0.879 | -0.16 |
|  | ~ Adolescence |  | 0.94 | 0.33 | 0.005 | 1.17 |
|  | ~ Childhood +<br>adolescence |  | -0.29<br>0.99 | 0.37<br>0.34 | 0.44<br>0.004 | 1.11 |
| AUDIT at adolescence |  | 438 |  |  |  |  |
|  | ~ Birth |  | 0.70 | 0.38 | 0.064 | 0.56 |
|  | ~ Adolescence |  | 0.93 | 0.40 | 0.021 | 0.99 |
|  | ~ Birth +<br>adolescence |  | 0.56<br>0.83 | 0.38<br>0.41 | 0.138<br>0.042 | 1.27 |
| AUDIT at midlife |  | 518 |  |  |  |  |
|  | ~ Pregnancy |  | 0.15 | 0.26 | 0.557 | -0.13 |
|  | ~ Midlife |  | 1.63 | 0.25 | <0.001 | 7.38 |
|  | ~ Pregnancy +<br>midlife |  | -0.14<br>1.66 | 0.25<br>0.26 | 0.574<br><0.001 | 7.25 |

For ARIES children, we were able to compare the cross-sectional DNAm-Alc/AUDIT relationship in adolescence to DNAm-Alc measured at two earlier no/low drinking time points in the same individuals: in childhood (n=601) and at birth (n=438). AUDIT in adolescence appeared to be independent of DNAm-Alc in childhood (p = 0.88), including after adjusting for adolescent DNAm-Alc (p = 0.44). However, the cross-sectional relationship between DNAm-Alc and AUDIT in adolescence, while small in total magnitude (adjusted R^2^ = 1.2), persisted with similar effect size when adjusting for childhood DNAm-Alc (β = 0.94 compared to β = 0.99). While DNAm-Alc at birth showed some suggestive evidence of a relationship with AUDIT in adolescence when considered alone (β =0.70, p = 0.06), this effect attenuated when adjusted for cross-sectional DNAm-Alc (β =0.56, p = 0.14). We considered whether the suggestive relationship between DNAm-Alc levels at birth and AUDIT in adolescence could have been driven by maternal drinking during *in utero* development, but this did not appear to be the case in follow up analysis that added examination of maternal DNAm-Alc during pregnancy and which failed to show effects independent of DNAm-Alc at birth (Supplementary Table 5).

## Discussion

Using a population-based longitudinal study with repeated measures of alcohol use and blood DNA methylation in two generations and an additional independent replication population, we conducted in-depth validations of the DNAm-Alc biomarker scores developed by Liu et al., including their 144 CpG recommended DNAm-Alc score.^6^ Our results confirmed the utility of this biomarker as a measure for alcohol consumption in the general population, albeit with slightly decreased explanatory power than initially reported in adults without substantial levels of consumption. Performance was closer to initial assessment in our replication analysis using a population with much greater average levels of intake. Further, we demonstrated the DNAm-Alc score shows stronger correlation for sustained alcohol use compared to shorter term alcohol use, as evidenced by the higher proportion of the variance explained at midlife than in adolescence, and for AUDIT score compared to cross-sectional weekly alcohol intake. Additionally, by observing that an earlier measure of DNAm-Alc repeated in the same participants appears independent of later drinking behaviour, we show that this DNAm-derived score captures information reflective of alcohol-use exposure and behaviour, beyond genetic determinants alone (i.e. ruling out genetic confounding as the primary explanation of the results).

We observed stronger relationships between the DNAm-Alcs and AUDIT score, that captured additional behaviours related to alcohol dependency, compared to simple intake that the DNAm-Alcs were developed to predict. However, this may suggest that biological or behavioural factors reflecting individual adaptation to alcohol use are additionally being represented in the epigenetic response. This feature of the DNAm-Alcs may improve the utility of such scores in a clinical context, where alcohol dependency traits are likely more medically relevant than simple intake alone. However, we cannot exclude the possibility that the additional explanatory capacity we observed of DNAm-Alcs for alcohol dependency related features could also be linked to measurement error. For example, the dependency behaviour captured by the AUDIT measures may be more stable over time and thus more accurately capture long-term alcohol use, whereas cross-sectional self-report of alcohol intake is likely to be noisier due to the additional variability of drinking behaviour itself over time, as well as the measurement error introduced by trying to recall “average” recent drinking behaviour.

Compared to the variance of alcohol intake explained by the DNAm-Alcs in Liu et al’s initial report,^6^ we observed a fairly systematic reduction. This was expected, given the criticism described by Hattab, Clark and van den Oord^16^ that the evaluation of the DNAm-Alcs performed by Liu et al. suffered from statistical overfitting due to re-estimation of coefficient values in testing data. Thus, while the adjusted R^2^ of the 144CpG DNAm-Alc in the adult analyses in that study were seen to range from 12.0 to 13.8%, in our validation computed using the independently estimated Liu et al. coefficients in a comparable sample – ARIES parents at midlife - we observed roughly half the explanatory capacity of that score (Table 1, adjusted R^2^ = 7.63%). Further, when we reproduced the validation approach of Liu et al. in the same ARIES sample, we again observed performance metrics similar in magnitude to their initial findings (e.g. adjusted R^2^ of 12.47% for the 144CpG model in addition to the explanatory effect of age, sex and BMI), suggesting that the performance decrease we observed when applying the independently estimated coefficients to ARIES was due to the elimination of effects due to over-fitting. Importantly, we have facilitated future application of the correct, independently estimated Liu et al. DNAm-Alc coefficients through our freely distributed R software package, *dnamalci* (https://github.com/yousefi138/dnamalci).

While the DNAm-Alc showed limited explanatory capacity in ARIES, its performance was markedly improved in our HN5000 replication population, achieving 14% of variation explained (Supplementary Table 3). Participants in this study differed from our ARIES population in several important ways, including that they were all oropharyngeal cancer cases, slightly older, and had a little more than three times greater and more variable average alcohol intake. With the data available, we could not determine which factors were most responsible for the improved DNAm-Alc performance in HN5000, but it is at least possible that the biological impact of greater alcohol abuse in this group may be contributing.

We also observed reduced capacity of the DNAm-Alcs to predict categories of drinkers by alcohol intake in ARIES compared to the estimates by Liu et al. (Figure 3). In fact, the best performing DNAm-Alc in ARIES – AUC = 0.60 with the 144 CpG score for heavy versus non-drinkers – had a lower AUC than any DNAm-Alc for resolving heavy from light or non-drinkers at initial report. The lowest that had been observed there being AUC = 0.67 for heavy versus light using the 5 CpG score in the ARIC study. However, our ability to evaluate prediction performance was severely limited in power by the small number of participants falling into the higher categories of intake, with only a maximum of N=14 heavy drinkers for any of our comparisons. Interestingly, we did observe some ability of the DNAm-Alcs to resolve participants with different categories of AUDIT score, specifically high (>15) versus low AUDIT individuals, at midlife where we’d also seen the strongest relationship in continuous analysis. The two best-performing DNAm-Alcs in this regard, the 78 and 144 CpG biomarkers, had AUCs of 0.80 and 0.79 which is in a range comparable to that reported by Liu et al. for intake categories.

Our assessment of the timing of the relationship between alcohol use and the DNAm-Alcs, the third aim of our study, observed that these scores captured an epigenetic response largely independent of confounding from genetic or environmental predisposition to alcohol dependency. This was most clearly observed in the ARIES mothers, who had a strong relationship between DNAm-Alc and AUDIT when both were measured in participants at the same time but for whom almost no relationship was observed for DNAm-Alcs calculated from DNAm measured roughly 20 years previously, at a time when alcohol consumption was much lower during the women’s index pregnancy (Table 2). This was true both when the earlier DNAm-Alcs were considered on their own, or when mutually adjusted for the cross-sectional DNAm-Alc. If DNAm-Alc values were being driven largely by stable genotype-methylation relationships, as one might predict *a priori* given that DNAm levels are often highly correlated with nearby genetic polymorphisms,^17^ we would expect to observe a consistent relationship between such biomarker scores and alcohol dependency in the same participants regardless of the timing of the DNAm collection. However, this was not the case: when DNAm-Alcs were drawn at the earlier time point, no independent relationship was observed. We saw a similar trend among ARIES offspring in childhood and adolescence.

Further, our analysis of repeated measures of DNAm-Alcs in the same participants across time, and the comparisons of these across populations with different drinking behaviours and a different drinking history (young and middle-aged), helped identify the duration of exposure potentially reflected by these biomarkers. The DNAm-Alcs appeared to reflect more than just concurrent intake over a short-term window, like that captured by a breathalyser. The correlation observed between DNAm-Alcs and intake/AUDIT in ARIES adolescents with comparable levels of alcohol use to adults over a shorter time frame was much smaller than that of ARIES parents at midlife with longer histories of alcohol use (Table 1). This suggests that DNAm-Alcs may capture longer-term patterns of alcohol abuse, perhaps cumulatively (the ALSPAC parents having been using alcohol for over two decades, while their children are very recent users). It also suggests that the alcohol dependence and problem drinking components of the AUDIT scores could be driving these associations more than the actual alcohol consumption levels per se (alcohol behaviour in midlife being more likely to score highly on these components in British populations).^36^ However, this could not be evaluated specifically by our current study. While effects in adolescents were systematically smaller than for adults, they did appear to be more than just background noise: showing associations independent of the effects in those same participants at earlier time points with no likely alcohol exposure, at birth and 7 years of age (Table 2).

The results of our current study should be interpreted in the context of the following limitations: 1) Tissue specificity-DNA methylation measured in peripheral blood WBCs was used both for the original development of the DNAm-Alcs and our current validation. Blood may not be the tissue most directly relevant for assessing the impacts of alcohol abuse (as opposed to e.g. liver or brain). However, alcohol is absorbed by peripheral-blood and is relevant to the direct assessment of alcohol exposure. Further, blood-based biomarkers may be more clinically useful, less invasive, and more readily accessible than liver or brain tissue. 2) Measurement bias and measurement error-Cross-sectional measurement of alcohol use by self-report may be too crude an approximation of the biologically relevant exposure, due to the variability of alcohol behaviour over time. This has the potential to bias our estimates towards the null, particularly for alcohol intake as it is more variable than the AUD risk assessed by AUDIT. The latter includes a measure of, but is not limited to, alcohol consumption, but also reflects symptoms of dependence and indicators of alcohol-related problems. Previous reports have shown that AUDIT scores are more stable (than alcohol use reports) and particularly so in midlife compared to young adulthood, and this is probably attributable to a more variable nature of alcohol use in younger age.^37^ Further, variability in some individual’s alcohol use behaviour or reporting over time, combined with uncertainty about the temporality of alcohol-related changes in DNAm, could also contribute to measurement error. Such error could also potentially bias our results should it have occurred differentially. These mechanisms could play differing, possibly opposing roles at different points throughout the life course e.g. if under-reporting may be more common during pregnancy or true drinking behaviour may be especially variable during adolescence. 3) Residual confounding - We acknowledge the possibility of residual confounding in particular by unmeasured covariates acting in a time-dependent manner, i.e. exhibiting onset between the collection ages we have considered. 4) Statistical power - Although we had adequate power to perform most replications of the Liu et al. study, comparisons including heavy drinkers did suffer from low numbers of ARIES participants in this category. 5) Generalizability – Our study was limited to participants of European genetic ancestry, similar to those populations used by Liu et al. in their original development of the DNAm-Alcs. As such, our findings regarding DNAm-Alcs are not readily generalizable or valid for populations of other ancestries.

Our study also had notable strengths, including access to repeated DNAm measurements in participants drawn from a prospective, population-based, longitudinal birth cohort. These repeated observations, some of which were collected at time points prior to or during limited alcohol use, were unique in structure and allowed assessment of the role of genetic confounding that would not have been possible with many other study designs.

In conclusion, while we find that the DNAm-Alcs do not always predict alcohol intake as well as initially hoped, our revised evaluation in a large population-based sample demonstrates that they do still capture non-trivial amounts of information about the biological impacts related to alcohol use. Importantly, our results using AUDIT score rather than intake alone suggest that these biomarkers may explain additional variation in alcohol use disorder beyond alcohol consumption. Although DNAm signals often correlate strongly with local polymorphisms, our analysis in repeated measures from two separate generations was able to find strong evidence that DNAm-Alcs are largely independent of genetic confounding. The stronger relationship we observed between DNAm-Alcs and AUDIT in parents at midlife compared to offspring in adolescence despite similar levels of crosssectional consumption suggests that DNAm-Alc may reflect the biological impact of alcohol use over time. This was echoed in our findings from HN5000 which showed the strongest DNAm-Alc - intake relationship and was the population with the greatest amount of consumption. Overall, we confirm Liu et al.’s initial conclusion that DNAm-Alcs may have clinical utility as a test for detecting heavy alcohol consumption, but with some evidence that such markers track long term patterns of alcohol use rather than recent intake.

## Acknowledgements

The authors would like to thank the CHARGE alcohol working group for generously providing the coefficient effect estimates needed to replicate the DNAm-Alc analysis. Further, The authors would like to acknowledge the assistance of Michelle Taylor and Gemma Hammerton regarding the use of ALSPAC alcohol data, as well as helpful discussions with Jon Heron.

We are extremely grateful to all the families who took part in the ALSPAC study, the midwives for their help in recruiting them, and the whole ALSPAC team, which includes interviewers, computer and laboratory technicians, clerical workers, research scientists, volunteers, managers, receptionists and nurses.

We are also very grateful to all of those with head and neck cancer who took part in the Head and Neck 5000 study and would also like to thank the research, laboratory and clinical staff who supported this study.

## References

1 Rehm J, Mathers C, Popova S, Thavorncharoensap M, Teerawattananon Y, Patra J. Global burden of disease and injury and economic cost attributable to alcohol use and alcohol-use disorders. Lancet (London, England) 2009; 373: 2223–33.

2 De Bellis MD, Keshavan MS, Beers SR, Hall J, Frustaci K, Masalehdan A et al. Sex differences in brain maturation during childhood and adolescence. Cereb Cortex 2001; 11: 552–557.

3 Sen P, Shah PP, Nativio R, Berger SL, Adams PD, Armstrong VL et al. Epigenetic Mechanisms of Longevity and Aging. Cell 2016; 166: 822–839.

4 Tu W, Chu C, Li S, Liangpunsakul S. Development and validation of a composite score for excessive alcohol use screening. J Investig Med 2016; 64: 1006–1011.

5 Joehanes R, Just AC, Marioni RE, Pilling LC, Reynolds LM, Mandaviya PR et al. Epigenetic Signatures of Cigarette Smoking. Circ Cardiovasc Genet 2016;: CIRCGENETICS.116.001506.

6 Liu C, Marioni RE, Hedman ÅK, Pfeiffer L, Tsai P-C, Reynolds LM et al. A DNA methylation biomarker of alcohol consumption. Mol Psychiatry 2016. doi:10.1038/mp.2016.192.

7 Wahl S, Drong A, Lehne B, Loh M, Scott WR, Kunze S et al. Epigenome-wide association study of body mass index, and the adverse outcomes of adiposity. Nature 2016. doi:10.1038/nature20784.

8 Richard MA, Huan T, Ligthart S, Gondalia R, Jhun MA, Brody JA et al. DNA Methylation Analysis Identifies Loci for Blood Pressure Regulation. Am J Hum Genet 2017; 101: 888–902.

9 Bohlin J, Håberg SE, Magnus P, Reese SE, Gjessing HK, Magnus MC et al. Prediction of gestational age based on genome-wide differentially methylated regions. Genome Biol 2016; 17: 207.

10 Elliott HR, Tillin T, McArdle WL, Ho K, Duggirala A, Frayling TM et al. Differences in smoking associated DNA methylation patterns in South Asians and Europeans. Clin Epigenetics 2014; 6: 4.

11 Bojesen SE, Timpson N, Relton C, Davey Smith G, Nordestgaard BG. AHRR (cg05575921) hypomethylation marks smoking behaviour, morbidity and mortality. Thorax 2017;: thoraxjnl-2016-208789.

12 Vadigepalli R, Hoek JB. Introduction to the Virtual Issue Alcohol and Epigenetic Regulation: Do the Products of Alcohol Metabolism Drive Epigenetic Control of Gene Expression in Alcohol-Related Disorders? Alcohol Clin Exp Res 2018; 42: 845–848.

13 Joubert BR, Felix JF, Yousefi P, Bakulski KM, Ligthart S, Wang T et al. DNA Methylation in Newborns and Maternal Smoking in Pregnancy: Genome-wide Consortium Metaanalysis. Am J Hum Genet 2016; 98: 680–696.

14 Richmond R, Suderman M, Langdon R, Relton C, Smith GD. DNA Methylation As A Marker For Prenatal Smoke Exposure In Adults. bioRxiv 2017;: 121558.

15 Psaty BM, O’Donnell CJ, Gudnason V, Lunetta KL, Folsom AR, Rotter JI et al. Cohorts for Heart and Aging Research in Genomic Epidemiology (CHARGE) Consortium: Design of Prospective Meta-Analyses of Genome-Wide Association Studies From 5 Cohorts. Circ Cardiovasc Genet 2009; 2: 73–80.

16 Hattab MW, Clark SL, van den Oord EJCG. Overestimation of the classification accuracy of a biomarker for assessing heavy alcohol use. Mol Psychiatry 2017. doi:10.1038/mp.2017.181.

17 Gaunt TR, Shihab HA, Hemani G, Min JL, Woodward G, Lyttleton O et al. Systematic identification of genetic influences on methylation across the human life course. Genome Biol 2016; 17: 61.

18 Sanchez-Roige S, Fontanillas P, Elson SL, Gray JC, de Wit H, Davis LK et al. Genome-wide association study of alcohol use disorder identification test (AUDIT) scores in 20 328 research participants of European ancestry. Addict Biol 2017. doi:10.1111/adb.12574.

19 Saunders JB, Aasland OG, Babor TF, de la Fuente JR, Grant M. Development of the Alcohol Use Disorders Identification Test (AUDIT): WHO Collaborative Project on Early Detection of Persons with Harmful Alcohol Consumption--II. Addiction 1993; 88: 791–804.

20 Boyd A, Golding J, Macleod J, Lawlor DA, Fraser A, Henderson J et al. Cohort Profile: the ‘children of the 90s’--the index offspring of the Avon Longitudinal Study of Parents and Children. Int J Epidemiol 2013; 42: 111–127.

21 Fraser A, Macdonald-Wallis C, Tilling K, Boyd A, Golding J, Davey Smith G et al. Cohort Profile: the Avon Longitudinal Study of Parents and Children: ALSPAC mothers cohort. Int J Epidemiol 2013; 42: 97–110.

22 Relton CL, Gaunt T, McArdle W, Ho K, Duggirala A, Shihab H et al. Data Resource Profile: Accessible Resource for Integrated Epigenomic Studies (ARIES). Int J Epidemiol 2015; 44:1181–1190.

23 Ness AR, Waylen A, Hurley K, Jeffreys M, Penfold C, Pring M et al. Establishing a large prospective clinical cohort in people with head and neck cancer as a biomedical resource: head and neck 5000. BMC Cancer 2014; 14.

24 Ness AR, Waylen A, Hurley K, Jeffreys M, Penfold C, Pring M et al. Recruitment, response rates and characteristics of 5511 people enrolled in a prospective clinical cohort study: head and neck 5000. Clin Otolaryngol 2016; 41: 804–809.

25 Organization WH. International Statistical Classification of Diseases and Related Health Problems 10th Revision. 2016. http://www.who.int/classifications/icd/en/.

26 Deschler MG; Smith, RV DM. Quick Reference Guide to TNM Staging of Head and Neck Cancer and Neck Dissection Classification. 2014.

27 Saunders JB, Aasland OG, World Health Organization. WHO Collaborative Project on the Identification and Treatment of Persons with Harmful Alcohol Consumption. Report on phase I: the development of a screening instrument. Geneva, 1987.

28 Dedeurwaerder S, Defrance M, Calonne E, Denis H, Sotiriou C, Fuks F. Evaluation of the Infinium Methylation 450K technology. Epigenomics 2011; 3: 771–784.

29 Min JL, Hemani G, Davey Smith G, Relton C, Suderman M. Meffil: efficient normalization and analysis of very large DNA methylation datasets. Bioinformatics 2018;: bty476.

30 Fortin J-PP, Labbe A, Lemire M, Zanke BW, Hudson TJ, Fertig EJ et al. Functional normalization of 450 k methylation array data improves replication in large cancer studies. Genome Biol 2014; 15: 503.

31 Houseman EA, Accomando WP, Koestler DC, Christensen BC, Marsit CJ, Nelson HH et al. DNA methylation arrays as surrogate measures of cell mixture distribution. BMC Bioinformatics 2012; 13. doi:10.1186/1471-2105-13-86.

32 Jaffe AE, Irizarry RA. Accounting for cellular heterogeneity is critical in epigenome-wide association studies. Genome Biol 2014; 15: R31.

33 Bakulski KM, Feinberg JI, Andrews S V., Yang J, Brown S, L. McKenney S et al. DNA methylation of cord blood cell types: Applications for mixed cell birth studies. Epigenetics 2016; 11: 354–362.

34 Robin X, Turck N, Hainard A, Tiberti N, Lisacek F, Sanchez JJ-C et al. pROC: an open-source package for R and S+ to analyze and compare ROC curves. BMC Bioinformatics 2011; 12: 77.

35 R Core Team. R: A Language and Environment for Statistical Computing. 2018. https://www.r-project.org/.

36 Britton A, Ben-Shlomo Y, Benzeval M, Kuh D, Bell S. Life course trajectories of alcohol consumption in the United Kingdom using longitudinal data from nine cohort studies. BMC Med 2015; 13: 47.

37 Sahker E, Lancianese DA, Arndt S. Stability of the alcohol use disorders identification test in practical service settings. Subst Abuse Rehabil 2017; Volume 8: 1–8.

