## Supplementary information for "Validation and characterization of a DNA methylation alcohol biomarker across the life course"

| | | N | $R^2$ | | | |
|---|---|---|---|---|---|---|
| | | | 5 CpGs | 23 CpGs | 78 CpGs | 144 CpGs |
| Midlife | | | | | | |
| | Alcohol intake | 1 049 | 3.84 | 4.32 | 6.7 | 7.57 |
| | AUDIT | 1 013 | 5.04 | 5.73 | 8.85 | 9.00 |
| Adolescence | | | | | | |
| | Alcohol intake | 626 | 0.12 | 0.26 | 0.57 | 0.61 |
| | AUDIT | 620 | 0.00 | 0.05 | 0.66 | 0.88 |

Supplementary table 1. $R^2$ between DNAm-Alcs and measures of alcohol intake (log(g/day + 1)) and AUDIT score, in ARIES parents at midlife and offspring at adolescence with *adjustment of DNAm-Alcs for concurrent smoking*.

| | | N | $R^2$ | | | |
|---|---|---|---|---|---|---|
| | | | 5 CpGs | 23 CpGs | 78 CpGs | 144 CpGs |
| Midlife | | | | | | |
| | Alcohol intake | 988 | 3.80 | 4.28 | 6.89 | 7.58 |
| | AUDIT | 952 | 5.51 | 6.27 | 10.07 | 9.77 |
| Adolescence | | | | | | |
| | Alcohol intake | 586 | 0.13 | 0.35 | 0.57 | 0.64 |
| | AUDIT | 580 | 0.00 | 0.07 | 0.77 | 1.05 |

Supplementary table 2. $R^2$ between DNAm-Alcs and alcohol intake (log(g/day + 1)) in ARIES parents at midlife and offspring at adolescence, *excluding self-reported non-drinkers*.

| | | N | $R^2$ | | | |
|---|---|---|---|---|---|---|
| | | | 5 CpGs | 23 CpGs | 78 CpGs | 144 CpGs |
| HN5000 | | | | | | |
| | Alcohol intake | 281 | 8.88 | 7.34 | 12.52 | 14.34 |

Supplementary table 3. $R^2$ between DNAm-Alcs and alcohol intake (log(g/day + 1)) and AUDIT score in HN5000.

| | | N | Adjusted $R^2$* | | | | |
|---|---|---|---|---|---|---|---|
| | | | ~ Age + Sex + BMI | + 5 CpGs | + 23 CpGs | + 78 CpGs | + 144 CpGs |
| Midlife | | | | | | | |
| | Alcohol intake | 1 049 | 4.78 | 8.77 | 9.73 | 12.69 | 17.25 |
| Adolescence | | | | | | | |
| | Alcohol intake | 626 | 0.00 | -0.63 | -0.17 | -2.63 | -0.45 |

Supplementary Table 4. Adjusted $R^2$ from models regressing measures of alcohol use, alcohol intake (g/d), in ARIES parents at midlife and offspring at adolescence on the methylation β-values of CpGs used in constructing DNAm-Alcs, as was the modelling approach presented in Liu et al. 2016. *See Methods for details.

| Dependent Variable | Independent Variable(s) (DNAm Alc score) | N | β | SE | P-value | Adjusted $R^2$* |
|---|---|---|---|---|---|---|
| AUDIT at adolescence | | 380 | | | | |
| | ~ Birth | | 0.70 | 0.40 | 0.08 | 0.55 |
| | ~ Pregnancy | | 0.46 | 0.43 | 0.29 | 0.04 |
| | ~ Birth + | | 0.67 | 0.40 | 0.10 | 0.49 |
| | Pregnancy | | 0.38 | 0.43 | 0.38 | |

Supplementary Table 5. Estimates of the effects of DNAm-Alc in ARIES offspring at birth and in mothers during pregnancy on offspring AUDIT at adolescence considered both in separate single-predictor models and simultaneously.
